# An open, integrated platform for multiplexed bioluminescence microscopy

**DOI:** 10.1101/2025.08.17.670750

**Authors:** Lorenzo Scipioni, Belen Torrado, Giulia Tedeschi, Lila P. Halbers, Zachary R. Torrey, Erin B. Fuller, Francesco Fersini, Christoph Gohlke, Andrej Luptak, Jennifer A. Prescher, Michelle A. Digman

**Affiliations:** University of California, Irvine (CA, United States), Department of Biomedical Engineering, Laboratory for Fluorescence Dynamics; Centre de Recherche en Cancérologie de Toulouse, Toulouse (France); University of California, Irvine (CA, United States), Beckman Laser Institute and Medical Clinic; University of California, Irvine (CA, United States), Department of Pharmaceutical Sciences; University of California, Irvine (CA, United States), Department of Chemistry; University of California, Irvine (CA, United States), Department of Molecular Biology & Biochemistry

## Abstract

We report an imaging package to democratize all-in-one bioluminescence and fluorescence microscopy. The platform comprises three tools: PhasorViewer, a visualization suite to design experiments and identify optimal probe combinations; PhasorScope, an open-source, cost-effective microscopy framework to upgrade conventional microscopes; and PhasorAnalysis, a dedicated, user-friendly analysis pipeline to quantify phasor imaging data. We demonstrate the utility of the platform for multiplexed, simultaneous fluorescence and bioluminescence imaging with readily accessible optical reporters.

## Main text

Imaging tools provide “eyes” into biology and fuel new discoveries.^1-3^ There remains a gap in our ability to see dynamic processes at the single-cell level, though, especially in heterogeneous environments. Conventional fluorescence approaches rely on excitation light that can induce photobleaching and phototoxicity, hindering many pursuits.^4-5^ Bioluminescent platforms can circumvent these issues,^2,6-10^ and dozens of luciferases are now available for microscopy applications.^11-14^ However, their spectra are broad and overlapping, complicating multi-component studies^9,15-16^ Imaging with more than one reporter—commonplace in fluorescence—is incredibly rare in bioluminescence microscopy.^17^

Bioluminescent phasor addresses the key gap in multiplexing ability. This approach optically transforms spectral signatures into spectral phasors, providing phasor coordinates with high spectral resolution and uncompromised integration times.^18^ Bioluminescent phasor also retains on average twice as many photons as RGB cameras.^20^ We previously showcased the utility of bioluminescent phasor for multicolor longitudinal imaging of both subcellular features and heterogeneous spheroids. We further resolved previously indistinguishable mixtures of optical reporters, and provided universal and precise measurements of bioluminescence resonant energy transfer (BRET), a parameter of crucial importance in biosensor design^19^.

While we demonstrated feasibility, key hurdles prevented easy adoption of bioluminescent phasor technology. The instrument relied on two cameras and thus a complicated calibration scheme. The initial design also did not support fluorescence and lacked a user-friendly analysis software to resolve optical reporters. Furthermore, identifying good combinations of reporters for multiplexed imaging *a priori* was nontrivial.

Here we report a compact, user-friendly workflow that overcomes these obstacles and makes multiplexed bioluminescence and fluorescence imaging accessible to the broader community (**Figure 1**). First, we developed PhasorViewer, a simple, interactive tool for identifying combinations of bioluminescent probes for microscopic imaging. Second, we designed PhasorScope, an open-microscopy platform that is easily integrated with conventional microscopes. A complete parts list (totaling ∼$7500 in 2025, excluding the price of the camera) and manual, along with a step-by-step video, guide the user through the assembly, alignment, and calibration. The PhasorScope build requires only ∼2 h for an experienced engineer, and ∼8 h for a non-specialist. Each subsequent use only requires alignment and calibration checks that are performed in minutes. Last, we produced PhasorAnalysis, an analysis pipeline featuring cell segmentation and a variety of common plotting and output files (e.g., spreadsheets and image files). PhasorAnalysis also features a simple user interface and cursors for real-time decisions, and is accompanied by a video tutorial on the code. PhasorViewer and PhasorAnalysis are written in Google CoLab, a browser-based platform for Python programming, to maximize access.

**Figure 1:**
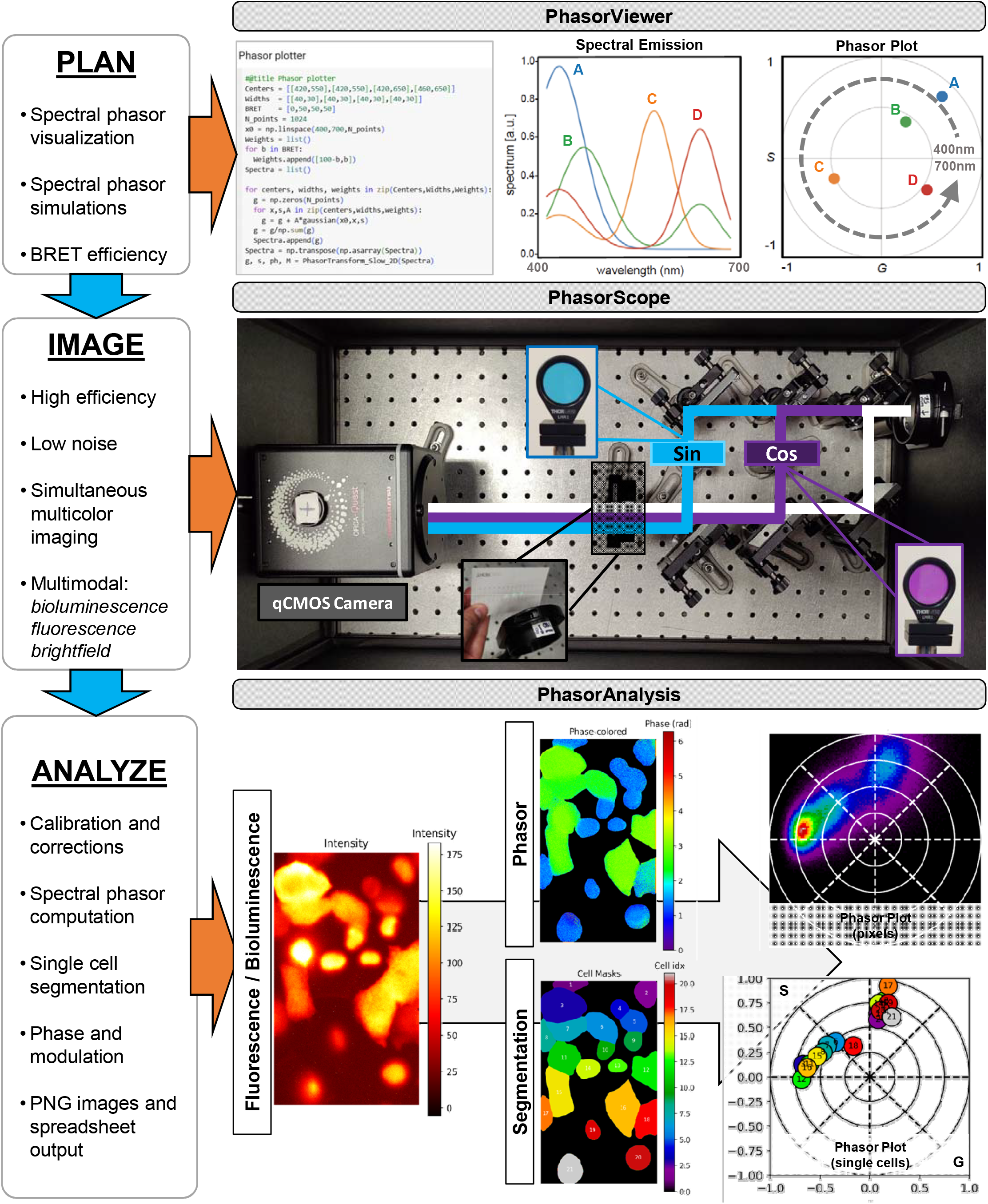
Workflow for integrated optical imaging. PhasorViewer: screenshot of Colab code (left), plot of simulated complex spectra (center) and corresponding phasor coordinates (right) for selecting optical reporters. **PhasorScope**: photo of the assembly, highlighting the optical paths for unfiltered (white), sine-filtered (cyan) and cosine-filtered (purple) light. Insets show pictures of the sine and cosine filters as well as the three optical paths. **PhasorAnalysis**: sample output images (Intensity, Phasor and Segmentation) and plots (pixels and single-cells). Numbers and colors in the single-cell phasor plot correspond to the ones in the segmentation image.

To showcase the modularity of PhasorScope, we used two commercial cameras, the Hamastsu Orca Quest and the Nüvü HNü 512 featuring different technologies (qCMOS and EMCCD), pixel sizes (4.6 μm and 16 μm) and numbers (4096×2304 and 512×512, **Figure 2A**). Both setups could readily detect cells expressing bioluminescent enzymes (YeNL in Figure 2A, NanoLuc and LumiScarlet in Figure S1). The analysis platform also readily reported similar phasor locations for each reporter. The system is thus robust and amenable to changes in components. Moreover, the spectral phasor coordinates can be directly compared across instruments and laboratories.

**Figure 2.**
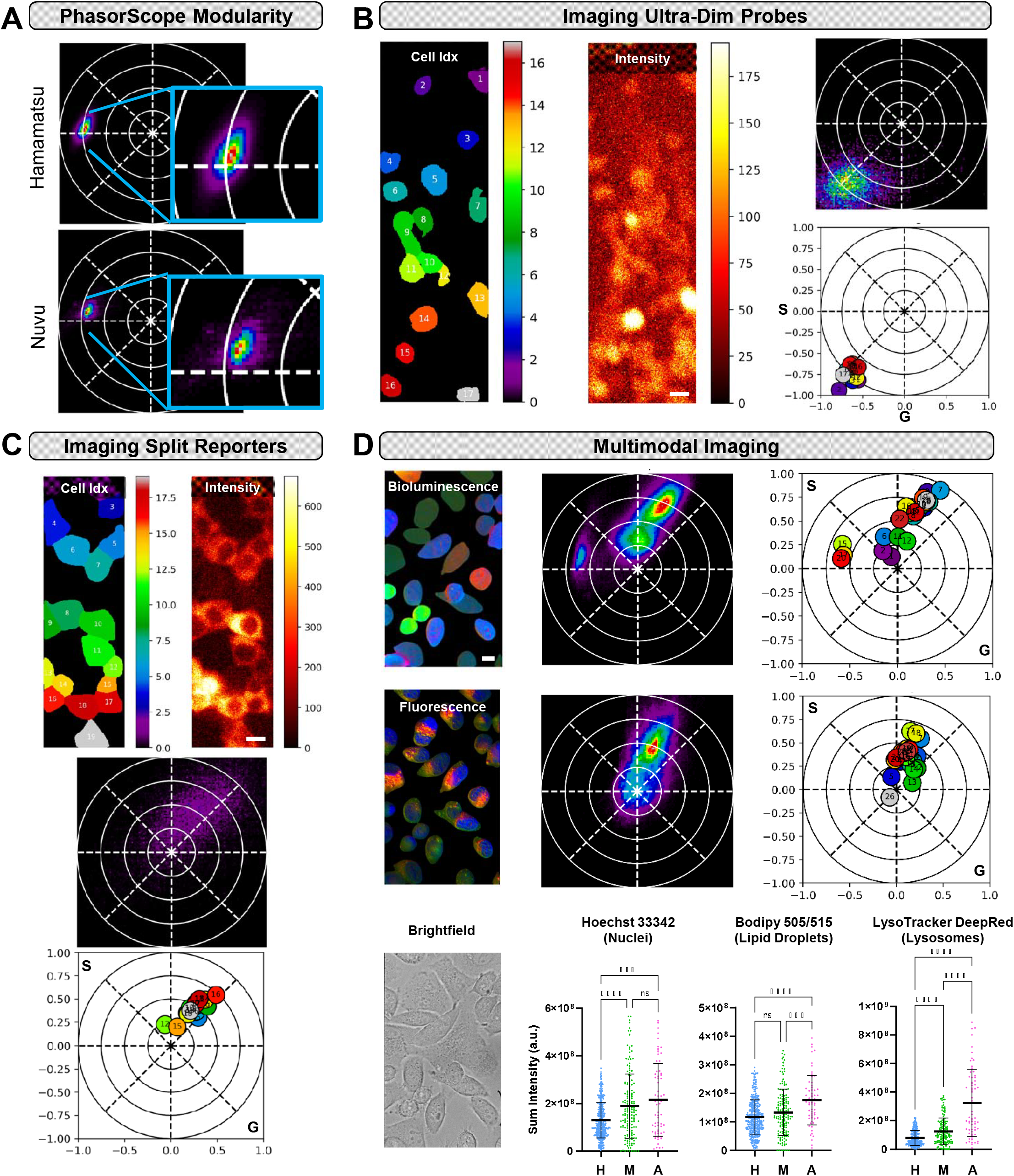
Applications of PhasorScope. **(A)** PhasorScope imaging with different cameras. MDA-MB231 cells stably expressing YeNL were imaged with a Hamamatsu qCMOS camera (left) or a Nüvü EMCCD camera (right). (**B**) Detection of dim-emitting probes. Fluc-expressing HEK cells were treated with D-lucifeirn and imaged. (**C**) RNA imaging with a split luciferase reporter. For (B)-(C), Segmented cells (left), the intensity image (middle), the pixel-wise phasor distribution (top right), and average phasor location of the single cells (bottom right) are shown. Scale bars = 10 µm. (**D**) Multimodal imaging (bioluminescence, fluorescence and brigthfield) with co-cultures of three cell lines. Three distinct live-cell organelle stains were also used. **Top row**, left to right: intensity-weighted unmixed bioluminescence image (HeLa BreakFAST Cyto-Nluc in blue, MDA-MB231 YeNL in green, A549 LumiScarlet in red), pixel-wise phasor plot of bioluminescence signal, average phasor location of the single cells for bioluminescence image. **Middle row**, left to right: intensity-weighted fluorescence image (nuclei in blue, lipid droplets in green, lysosomes in red), pixel-wise phasor plot of fluorescence signal, average phasor location of the single cells for fluorescence image. Bottom row, left to right: brightfield image of the same field of view as bioluminescence and fluorescence images. Boxplots show sum intensities distribution for nuclear (Hoechst 33342), lipid droplet (Bodipy 505/515) and lysosomal (LysoTracker DeepRed) signals, respectively, for each cell line (H= HeLa BreakFAST Cyto-Nluc, M = MDA-MB231 YeNL, A = A549 Lumiscarlet). Scale bar = 10 µm.

We further showcased PhasorScope for imaging ultra-dim signals. Firefly luciferase (Fluc)-expressing cells and D-luciferin are widely used for imaging in vivo, but the slow turnover rate has precluded many microscopic applications. More sensitive cameras are beginning to address the need, and we showed that one (Nüvü) was compatible with the PhasorScope. As shown in **Figure 2B**, Fluc-expressing cells were readily visible on the setup and registered phasors. Note that optical phasor transformation can result in phasors outside the unit circle, particularly in the presence of noise and narrow emission spectra. In a second example, we imaged cells expressing a split luciferase capable of RNA detection (RNA lantern, **Figure 2C**).^21^ These and other split reporters are notoriously dimmer compared to their intact counterparts, but our work shows that they amenable to bioluminescent phasor.

PhasorScope readily supports both fluorescence and bioluminescence imaging, a feature long-desired in the imaging community. As shown in **Figure 2D**, we sequentially acquired brightfield (using an illumination lamp), bioluminescence (with luciferin application), and fluorescence (LED excitation) images of live cells in the same field of view. Three distinct cell types, each expressing a different luciferase (NanoLuc, LumiScarlet, or YeNL), were observed, along with three unique three organelle-targeting fluorophores. PhasorScope imaging provides cell morphology (brightfield), cell type (bioluminescence) and physiological signature (organelle dyes) at optical resolution in the same field of view. We then used PhasorAnalysis on the bioluminescence and fluorescence images to quantify the intensity of each luciferase and organelle dye. Ultimately, we were able to simultaneously quantify the properties for the three organelles in all three cell lines from a single, multimodal dataset.

In conclusion, we developed an open-source, open-microscopy toolbox for planning (PhasorViewer), imaging (PhasorScope) and analysis (PhasorAnalysis) with uncompromised optical resolution and integration, democratizing multiplexed bioluminescence and fluorescence imaging for the research community.

## Acknowledgments

This work was supported by the U.S. National Institutes of Health (P41-GM103540 to L.S. and M.A.D.), the Paul G. Allen Frontiers Group (to J.A.P and M.A.D), and the Chan Zuckerberg Initiative (to J.A.P. and M.A.D.). L.S., G.T. and F.F. were also supported by the Chaire Oncobreast, Fondation Toulouse Cancer Santé. We thank Prof. Enrico Gratton and members of the Laboratory of Fluorescence Dynamics (LFD, UCI) for helpful discussions.

## Author contributions

L.S., B.T., G.T., M.A.D. and J.A.P. conceived the project idea. L.S., B.T., G.T., L.P.H., Z.R.T., and E.B.F. performed the experiments. L.S. and B.T. designed and built the imaging setup. L.S., F.F., and C.G. wrote the codes and analyzed the raw imaging data. A.L. provided technical support on the imaging setup. L.S., B.T., G.T., L.P.H., Z.R.T., E.B.F., M.A.D., and J.A.P. analyzed data and contributed to the writing of the manuscript. All authors have given approval to the final version of the manuscript.

## Materials and Methods

### General cell culture methods

HEK293, MDA-MB-231, HeLa, and A549 cells (ATCC), and cells engineered to express various reporters were cultured in complete media: DMEM (Corning) containing 10% (v/v) fetal bovine serum (FBS, Life Technologies), 4.5 g/L glucose, 2 mM L-glutamine, penicillin (100 U/mL), and streptomycin (100 µg/mL, Gibco). For transient transfection experiments, HEK293 cells (5□×□10^5^) were plated 24–48 h prior to transfection in tissue culture treated 6-well dishes (Corning). Transfections were performed with luc2-IRES-eGFP (Fluc) or Staygold-MS2-3-PP7 (RNA bait for RNA probe) using Lipofectamine 3000 according to the manufacturer’s instructions when cells were 75-80% confluent (1-2 d post plating). Cells were manipulated 24-48 h post transfection. Cells were incubated at 37 °C in a 5% CO2 humidified chamber. Cells were serially passaged using trypsin (0.25% in HBSS, Gibco).

### Viral transductions

FLAG-tagged YeNL and LumiScarlet reporter plasmids (in pLenti backbones), described in Table S1 were co-transfected with the VSV-G envelope plasmid and d8.2 gag/pol helper plasmids into Lenti X-293T cells (seeded in 10 cm dishes). Transfections were performed using 1 mg/mL polyethyleneimine (PEI, VWR #BT129700). A total of 1.5 μg VSV-G, 5 μg d8.2, and 6 μg FLAG-pLenti plasmids were mixed in 500 μL OptiMEM (LifeTech #1158021) and vortexed. In a separate tube, 25 μL PEI (2 μL PEI/μg DNA) was added to 475 μL OptiMEM and vortexed. Both solutions were incubated at room temperature for 5 min, then combined and incubated for an additional 20 min. The resulting mixture was added dropwise to the seeded Lenti X-293T cells. The following day, the cell culture medium was replaced with DMEM supplemented with 1x NaPyr, 10 mM HEPES, 1X GlutaMAX, and 1X PSG (no FBS). Two days later, the media was collected, filtered through a 0.45 μm PVDF membrane, and the viral particles were concentrated by ultracentrifugation (100,000 ×g for 2 h at 4 °C using a SW28 rotor on a Beckman L8-80M centrifuge) into a sucrose cushion. The concentrated virus was resuspended in cold PBS and either stored at –80 °C or used immediately on target cells (MDA-MB-231s and A549)

MDA-MB-231 Lumiscarlet, HeLa Nluc or MDA-MB231 YeNL were seeded in 8-well imaging chambers (Cellvis) and imaged with either the Hamamatsu (ORCA-Quest qCMOS, C15550, Hamamatsu) or the Nüvü (HNü 512, 512 × 512 EMCCD) camera. For each cell line, we acquired 2 frames with 2 minutes integration. Calibration data were acquired as described above and processed using the PhasorAnalysis code (Figure S1).qCMOS/EMCCD comparison experiment

**Table S1:** Table reporting on the name, features, vector type and reference to original manuscript and Addgene number for the constructs used in this work.

| Reporter | Features | Plasmid Vector | Reference | Addgene # |
| --- | --- | --- | --- | --- |
| NanoLuc | FLAG - Nluc - SV40 - PuroR - WPRE - AmpR | pLenti 3<sup>rd</sup> gen | Yao, Z.; et al. | 183045 |
| YeNL | FLAG - Venus - Nluc - SV40 - PuroR - WPRE - AmpR | pLenti 3<sup>rd</sup> gen | Yao, Z.; et al. | 208849 |
| LumiScarlet | FLAG - mScarlet - Nluc - SV40 - PuroR - WPRE - AmpR | pLenti 3<sup>rd</sup> gen | Yao, Z.; et al. | 208854 |
| RNA probe | SV40NLS- HA – MCP- SmBiT114- IRES- PCP- LgBiT- FLAG- PuroR - WPRE - AmpR | pLenti 3<sup>rd</sup> gen | Halbers, L. P.; et al. | 226095 |
| Staygold-M-3-P | Staygold - MS2-3-PP7- AmpR | pCDNA 3.1 | Halbers, L. P.; et al. | N/A |
| BreakFAST-cyto | BreakFAST- SV40 - PuroR - WPRE - AmpR | pLenti 3<sup>rd</sup> gen | Torrey, Z. R.; et al. | N/A |
| Firefly (Fluc) | luc2-IRES-eGFP- AmpR | pCDNA 3.1 | Love, A. C.; et al. | N/A |

### RNA probe, NanoLuc, and BreakFAST

Cell lines stably expressing RNA probe, NanoLuc, and BreakFAST were generated via lentiviral transduction. Briefly, HEK293 packaging cells were plated in 6-well dishes and transfected as described above with viral packaging plasmids along with luciferase- plasmids in the pLenti vector. Cells were transfected at 70% confluency. Twenty-four hours post transfection, media was replaced with viral induction media: DMEM (Corning) containing 4.5 g/L glucose, 2 mM L-glutamine, 20 mM HEPES buffer, and 10 mM sodium butyrate. After a 48 h incubation with viral induction media, media was collected from HEK cells and spun down at 500 x*g* for 5 min. Supernatant was plated directly on pre-plated and adhered destination lines (HEK, HeLa, MDA-MB-231s), 8 µg/mL polybrene (final concentration) was added and cells were incubated for 2 d. Media was then removed and fresh media added containing 25 µg/mL puromycin for selection. Final puromycin concentrations were tested and scaled for individual cell lines. Once selected cells grew to 100% confluency in a T25 or T75 flask, cells were subjected to FACS to select the brightest clones. Cells were seeded in 8-wells imaging chambers (CellVis) at a density of 30 000 cells/well, the day before imaging and kept in the incubator. Prior to imaging (10 min), 3 μL of furimazine (50 mM, Promega) was added.

### Fluc probe

HEK293 cells (5□×□10^5^) were seeded in 8-well Ibidi µ-Slides and transfected with Fluc-IRES-eGFP construct (Table S1) using Lipofectamine 3000 (ThermoFisher) following manufacturer instructions. Cells were imaged 24 h post-transfetion. Fluorescence illumination acquired using blue LED light source (ThorLabs, Solis-470C) of eGFP was used to localize positively expressing cells. For bioluminescence imaging data collection, it was added 3uL per well (total volume 300uL) of the substrate D-Luc from the stock 10 mM (Goldbio catalog #LUCK-100)

The following acquisition parameters were used: Nüvü camera objective 20x 0.75 NA, collection of 12 frames with a 5 min integration time per frame, 750 gain. Calibration was performed by acquiring “bright” (brightfield lamp, 20 frames, 200 ms integration time), “dark for bright” (no illumination, 20 frames, 200 ms integration time), “dark for measures” (12 frames, same integration time as the measures). Data were processed using the PhasorAnalysis Colab code.

### RNA Probe Imaging

HEK293 cells (5□×□10^5^) stably expressing the split RNA MS2 and PP7 coat protein lantern (RNA lantern) were plated in 8-well Ibidi µ-Slides. After 24□h, the cells were transiently transfected with *Staygold*-*MS2*-*3*-*PP7* (Table S1) using Lipofectamine 3000 (ThermoFisher) following manufacturer instructions. 24 hours after transfection cells were imaged. To carry on the measurement, we used the Phasor Scope with Nüvü camera for detection. To confirm the location of co-transfected cells we collected fluorescent measurements (LED laser 505nm emission, emission filter 535/30, 2 frames with 25ms integration time, gain 100). To collect the bioluminescence signal 2 frames with 2 min. integration time, gain 100 was used.

### Experiment for multimodal imaging (fluorescence, bioluminescence, brightfield)

For each experimental day, the following calibration datasets were collected:

⍰ Dark for fluorescence (20 frames, 200 ms integration time)
⍰ Dark for bioluminescence (2 frames, 2 minutes integration time)
⍰ Bright (brightfield lamp, intensity set to have ∼16k counts in detection, the acquired at 200 ms integration time, 20 frames).

For single cell lines with bioluminescence reporter, 3uL per well of substrate Furimazine (50 mM, Promega) was added 10 minutes before imaging. The focal plane was determined with brightfield illumination, acquiring 1 frame (200 ms integration time). Sequentially, we acquired a brightfield, a bioluminescence, a fluorescence, and again a brightfield image.

For co-culture experiments, a 1:1:1 mixture of HeLa BreakFAST Cyto-Nluc, A549 Lumiscarlet, MDA-MB231 YeNL were seeded in 8-well imaging chambers (Cellvis). The day of the experiment, culture media was replaced with media containing a mixture of organelle labeling dyes: Hoechst 33342 for DNA, BODIPY 505/515 for lipid droplets, Lysotracker DeepRed for lysosomes). Bodipy 505/515 (Cayman Chemicals, final concentration 1µM), LysoTracker DeepRed (ThermoFisher, final concentration 100 nM).

To calibrate the phasor positions for bioluminescence and fluorescence, we imaged single populations of HeLa BreakFAST Cyto-Nluc, A549 Lumiscarlet, MDA-MB231 YeNL alone in bioluminescence (Figure S2), and we stained HeLa Nluc cells with single dyes (fluorescence, Figure S3). Multimodal images were analyzed as follows. Bioluminescence images were used for cell segmentation (Cellpose), and the bioluminescence and fluorescence images were unmixed by spectral phasor unmixing. The cells were classified as HeLa, A549 or MDA-MB-231 according to which bioluminescent reporter (Cyto-Nluc, Lumiscarlet or YeNL, respectively) resulted in the highest relative fraction. For each classified cell line, the fluorescence signal of the organelle dyes was quantified as the total intensity of the unmixed signal within cell masks.

## Supplementary

**Figure S1.**
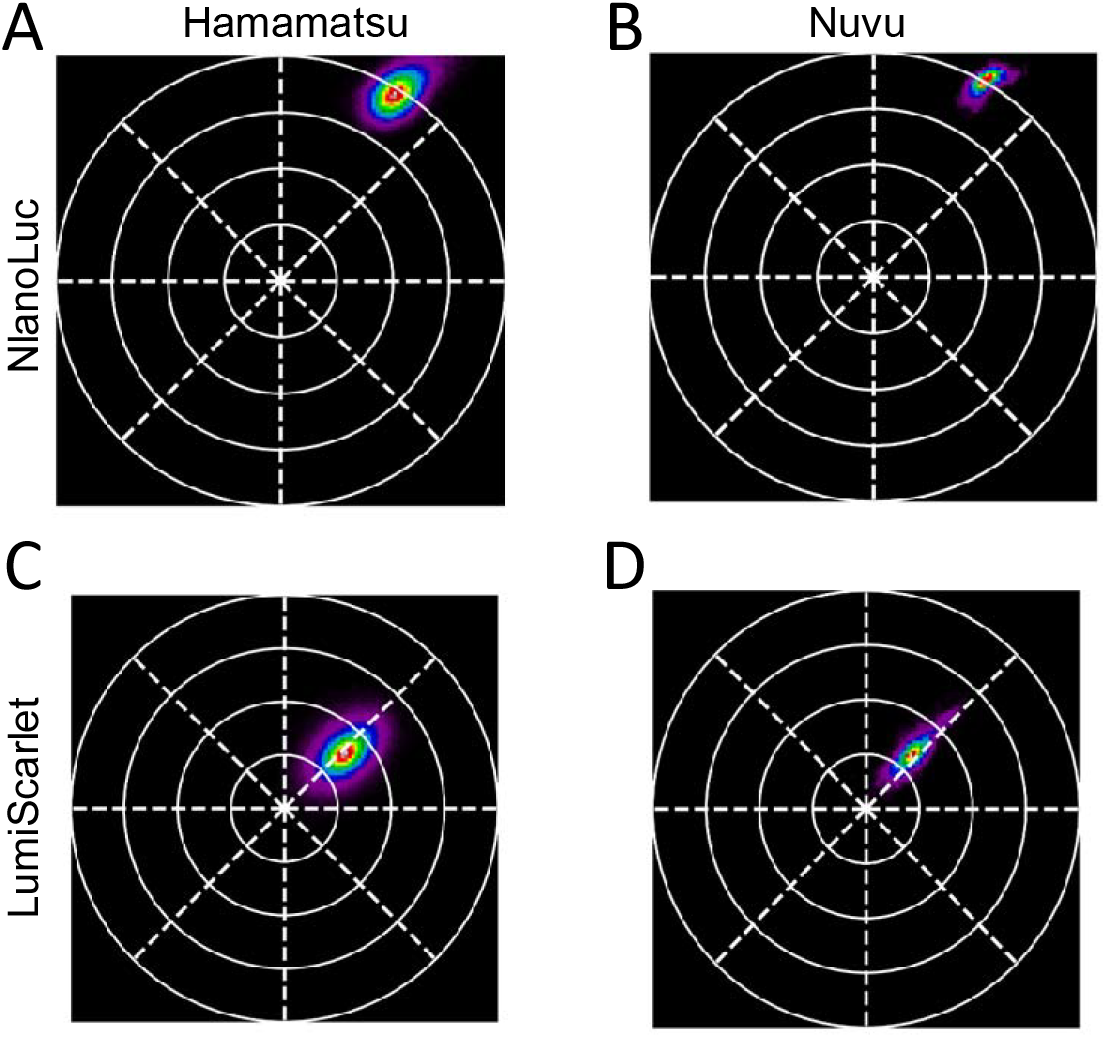
Bioluminescence reporters across cameras. MDA-MB231 cells stably expressing NanoLuc (**A**,**B**) or LumiScarlet (**C**,**D**) have been imaged in the PhasorScope equipped with the Hamamatsu camera (**A**,**C**) or the Nuvu camera (**B**,**D**).

**Figure S2.**
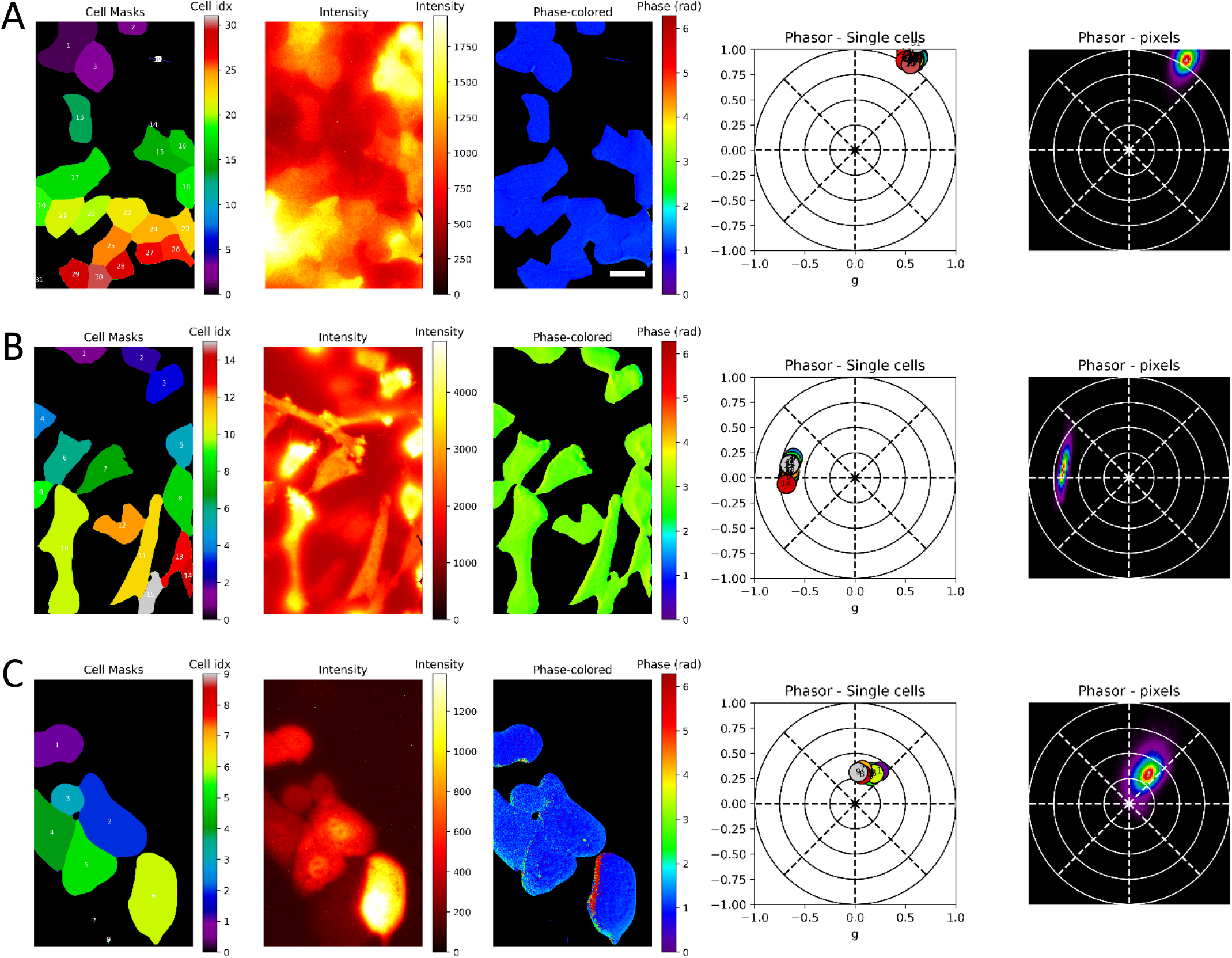
Bioluminescence phasor signature of 3 different report-expressing cell lines. Left to right: cell segmentation, intensity image, phasor phase colored image,, average phasor position per cell, pixel wise phasor distribution. (**A**) HeLa BreakFAST Cyto-Nluc, (**B**) MDA-MB231 YeNL, (**C**) A549 Lumiscarlet. Scale bar is 10 µm.

**Figure S3.**
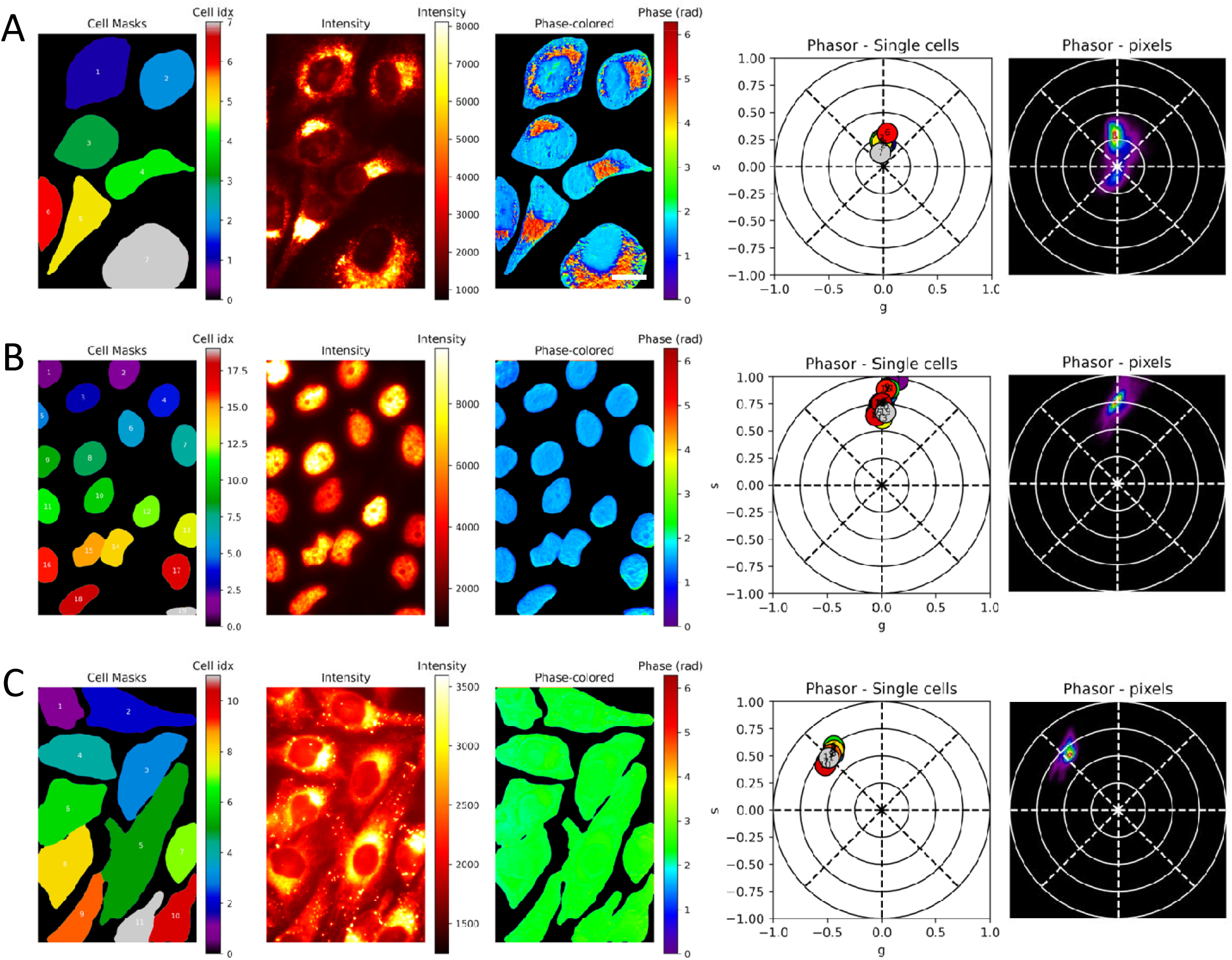
HeLa BreakFAST Cyto-Nluc labelled with live organelle dyes. Left to right: cell segmentation, intensity image, phasor phase colored image, average phasor position per cell, pixel wise phasor distribution. (**A**) LysoTracker DeepRed 100 nM, (**B**) Hoechst 33342 60nM, (**C**) Bodipy 505 1µM. Scale bar is 10 µm.

